# V3 Interneurons are Active and Recruit Spinal Motor Neurons During *In Vivo* Fictive Swimming in Larval Zebrafish

**DOI:** 10.1101/2021.03.03.433646

**Authors:** Timothy D. Wiggin, Jacob E. Montgomery, Amanda J. Brunick, Jack H. Peck, Mark A. Masino

## Abstract

Survival for vertebrate animals is dependent on the ability to successfully find food, locate a mate, and avoid predation. Each of these behaviors requires motor control, which is set by a combination of kinematic properties. For example, the frequency and amplitude of motor output combine in a multiplicative manner to determine features of locomotion such as distance traveled, speed, force (thrust), and vigor. Although there is a good understanding of how different populations of excitatory spinal interneurons establish locomotor frequency, there is a less thorough mechanistic understanding for how locomotor amplitude is established. Recent evidence indicates that locomotor amplitude is regulated in part by a subset of functionally and morphologically distinct V2a excitatory spinal interneurons (type II, non-bursting) in larval and adult zebrafish. Here we provide direct evidence, for the first time, that most V3 interneurons (V3-INs), which are a developmentally and genetically defined population of ventromedial glutamatergic spinal neurons, are active during fictive swimming. We also show that elimination of the spinal V3-IN population reduces the proportion of active MNs during fictive swimming but does not alter the range of locomotor frequencies produced. These data are consistent with V3-INs providing excitatory drive to spinal motor neurons and contributing to the production of locomotor amplitude, but not frequency, during swimming in larval zebrafish.

**SIGNIFICANCE STATEMENT:** Currently, there is a limited understanding about the cellular and spinal network properties that produce locomotor amplitude, defined as limb displacement in limbed animals or tail-bend in non-limbed animals during locomotion. Here we show, directly for the first time in a vertebrate, that V3 interneurons (V3-INs) in zebrafish larvae are active during *in vivo* fictive locomotion, and that targeted ablation of the spinal V3-IN population reduces the proportion of active motoneurons during fictive swimming. Importantly, ablation of V3-INs does not affect locomotor frequency (speed), which clarifies their role in motor control rather than rhythm generation. Thus, we propose that the V3-IN population is a source of excitation in the vertebrate locomotor neural circuitry that regulates locomotor amplitude independently of speed.

## INTRODUCTION

Efficient locomotion is set by a combination of kinematic properties, including the frequency and amplitude of rhythmic motor output. These properties combine to determine various locomotor features, such as distance traveled, speed, and vigor (Webb, 1975; Jayne and Lauder, 1995; Donley and Dickson, 2000; Tytell, 2004). Locomotor frequency (e.g. limb or tail-beat rhythmic activity) is a well-studied kinematic variable, which has been shown to increase linearly with speed (Bainbridge, 1958; Hunter and Zweifel, 1971; Webb et al., 1984; Müller and van Leeuwen, 2004). Less well studied is locomotor amplitude, which is measured by limb displacement in limbed animals during locomotor events (walking, trotting, galloping, hopping (Heglund et al., 1974)) or by tail-bend in non-limbed animals during anguilliform swimming (Wardle et al., 1995; Videler et al., 1999). Although there is a fundamental understanding of how identified subsets of excitatory spinal interneurons help to establish the frequency of motor output (Crone et al., 2009; Zhong et al., 2011; Ausborn et al., 2012; Eklöf-Ljunggren et al., 2012; Kimura et al., 2013; Ampatzis et al., 2014; Ljunggren et al., 2014; Menelaou et al., 2014; Bellardita and Kiehn, 2015), a more complete understanding of the cellular and network properties that regulate locomotor amplitude in vertebrate animals is lacking.

Across vertebrate species from fish to mammals, the neural circuitry that controls locomotion is located in the ventromedial spinal cord (Grillner and Jessell, 2009; Kiehn, 2011). In vertebrates, developmental gene expression specifies at least five cardinal classes of ventral neuronal progenitor cells. These progenitors give rise to motor neurons (MNs) and four classes of ventral interneurons, referred to as V0 to V3 neurons (Ericson et al., 1997; Pierani et al., 1999). Over the past decade, a population of identified excitatory spinal interneurons (V2a) has been shown to control the speed of locomotion in a variety of vertebrates (Zhong et al., 2011; Ausborn et al., 2012; Ampatzis et al., 2014; Gosgnach et al., 2017). More recent studies indicate a role for a subset of V2a interneurons (type II, non-bursting) in regulating locomotor amplitude (Menelaou and McLean, 2019) and/or vigor (Song et al., 2018) in larval and adult zebrafish, respectively. Both studies reveal a hierarchical control of motor neuron recruitment by a diverse V2a interneuron population, which the authors conclude is the basis for regulating locomotor amplitude and/or vigor.

To date, a cardinal class of spinal interneurons in the ventral cord that selectively regulates locomotor amplitude independently of locomotor frequency has not been identified. In the mammalian spinal cord, a genetically defined population of pre-motor glutamatergic neurons, called V3 interneurons (V3-INs), contributes to limb coordination, motor burst duration, and muscle spasms, but does not have a clear role in locomotor pattern generation (Chopek et al., 2018; Danner et al., 2019; Lin et al., 2019). We hypothesized V3-INs provide excitatory drive during locomotor activity to regulate locomotor amplitude. The work presented here shows directly, for the first time, that V3-INs in zebrafish larvae are active during *in vivo* fictive locomotion, and that elimination of the V3-IN spinal population reduces the proportion of active MNs during fictive swimming but does not alter the range of locomotor frequencies produced. These data are consistent with V3-INs providing excitatory drive to spinal motor neurons and contributing to the production of locomotor amplitude, but not frequency, during swimming in larval zebrafish. This is conceptually important because it clarifies a role of the V3 cardinal class of spinal interneurons in motor control, rather than rhythm generation.

## MATERIALS AND METHODS

### Fish Lines and Maintenance

All animal procedures were performed in accordance with the animal care committee’s regulations. Wild type (Segrest Farms, Gibsonton, FL) and transgenic adult zebrafish (*Danio rerio*) were maintained in the animal facility. The transgenic lines used in these experiments were: *Tg(nkx2.2a:mEGFP)^vu17^* (Kirby et al., 2006), *Tg(vglut2a:DsRed)^nns9^* (Miyasaka et al., 2009), *Tg(vglut2a:Gal4ff)^nns20^* (Satou et al., 2013), *Tg(UAS:GCaMP6s)^nk13a^* (Muto et al., 2017), and *parga^mn2Et^* (Balciunas et al., 2004). Out-crossed transgenic larvae at 4 – 6 days post fertilization (dpf) were used in these experiments. Larval zebrafish were maintained in petri dishes filled with embryo water (60 µg/ml Instant Ocean^®^ salt mix, Cincinnati, OH) and 0.00005% Methylene Blue in a 28.5°C incubator with a 14:10 light:dark cycle (ZT0 = 8:30am).

### V3-IN Soma Distribution Analysis

Quantification of the distribution of V3-IN cell bodies was performed using Fiji and custom Matlab code (Mathworks, Natick, MA). Confocal microscopy was used to collect overlapping stacks of images documenting the expression of *Tg(vglut2a:DsRed)^nns9^* and transmitted light morphology. Images were acquired at high magnification (40X / NA 0.8 objective) and a z-axis resolution of 1.3 µm. The fields of view in each stack overlapped with their neighbors by at least 10% of the width to permit accurate stitching and included the full width of the spinal cord and dorsal root ganglia (DRG) in the field of view. The collection of stacks was stitched into a single 3-D image of the caudal hindbrain and spinal cord (Preibisch et al., 2009). The Fiji Cell Counter tool (Schindelin et al., 2012)was used to identify the coordinates of all V3 neurons in the rostral 2 mm of the larval spinal cord (out of a total length of ∼3 mm), the coordinates of the DRG neuron clusters, and the position of the dorsal and ventral edges of the spinal cord. The markers tracing the dorsal and ventral edges of the spinal cord were used to calculate local tangent lines to the dorsal and ventral edges of the spinal cord at the rostrocaudal position of each V3-IN. The equation of a line perpendicular to the ventral tangent and intersecting the V3-IN was calculated, referred to as the “V3 plumb line.” The rostrocaudal distance of each V3 neuron was calculated as the distance along the ventral surface of the spinal cord between the spinal cord/hindbrain junction (Segment 2/3 boundary) and the V3 plumb line intersection with the ventral surface. The dorsoventral height of the spinal cord at the location of each V3-IN was calculated as the distance between the intersections of the V3 plumb line with the dorsal surface and the ventral surface, respectively. The distance of the V3-IN from the ventral surface of the spinal cord was also measured along the plumb line. The z-axis position of DRG neurons on each side of the spinal cord were used as boundary points of the lateral edges of the spinal cord, and a linear fit to the DRG neurons on each side of the animal was used to calculate the edges of the spinal cord at each rostrocaudal position in the imaging area. The midline slice and spinal cord width were calculated at the rostrocaudal position of each V3-IN, along with the distance from midline of the V3-IN.

### Whole-mount Immunohistochemistry

Five dpf *parga^mn2Et^* larvae were sacrificed with 0.2% Tricaine-S and fixed overnight in 4% paraformaldehyde at 4°C. Whole-mount larvae were permeabilized with Proteinase K, post-fixed, and blocked with 0.2% BSA, 10% normal donkey serum, 0.5% Triton X-100 in PBS. Goat polyclonal anti-choline acetyltransferase (ChAT; 1:50; AB144P) was added to the blocking solution and incubated overnight at 4°C. Fixed larvae were incubated with Alexa Fluor 633 donkey anti-goat secondary antibody (1:500; A-21082, Thermo Fisher Scientific Inc., Waltham, MA) in secondary blocking solution (0.2% BSA, 1% normal donkey serum, 0.5% Triton X-100 in PBS) for three nights at 4°C.

Anti-ChAT labeled *parga^mn2Et^* larvae were embedded laterally in low melting-point agarose. An Olympus Fluoview FV1000 confocal microscope was used to collect images through a 60x objective centered on body segment 15. Confocal stacks contained the entire dorsoventral extent of the spinal cord and spanned approximately three body segments (285 µm). Images were processed in Fiji and labeled cells from a single side of the spinal cord (approximately three hemisegments) were quantified using the Cell Counter plugin. Photoshop CS5 (Adobe Systems, San Jose, CA) was used to adjust brightness and contrast across entire z-projected images.

### In Vivo Rhod-2 AM Dye Loading of Motor Neurons

For *in vivo* characterization of MNs in *parga^mn2Et^* larvae, a 0.25% Rhod-2 AM (Life Technologies, Grand Island, NY) stock was made by dissolving 50 µg of dye in 20 µL Pluronic F-127:DMSO (1:4; Life Technologies, Grand Island, NY). Next, the skin was removed from the midbody region and a 1:50 dilution of the Rhod-2 AM stock was bath applied to the exposed muscles to permit sequestration of indicator dye by MNs via the peripheral motor axons. Sensory neurons were not strongly loaded by this procedure. Following one-hour incubation, the Rhod-2 AM solution was thoroughly washed-out using zebrafish ringers solution. Confocal images were collected, and cells were quantified as described for whole-mount immunohistochemistry (above).

### Peripheral Nerve Recordings and Analysis

Zebrafish larvae were anesthetized, the skin was carefully removed from the recording region, and larvae were paralyzed with of 0.1 mM α-bungarotoxin (Tocris, Ellisville, MO) to prevent muscle contractions. The anesthetic was removed to allow the larvae to perform fictive behavior, and a suction electrode (tip diameter: 9-15 µm) was placed over the intermyotomal cleft to record MN axons. Fictive motor bursts were identified and grouped into episodes using custom Matlab code. The program detected the presence or absence of activity at each voltage sample *v*(*n*). For each *v*(*n*), the algorithm determined a voltage autocorrelation *c_n_*(*k*) over a small window (3 ms) centered at *v*(*n*). These “windowed” autocorrelations were computed as *c_n_*(*k*) = 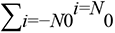 *v*(*n* − *i*)*v*(*n* − *i* − *k*) (*Eq. 1*), where 3-ms windowing was implemented in *Eq. 1* by setting *N*_0_ = (3 ms·*f*_sam_)/2, where *f*_sam_ is the sampling frequency, and by setting *v*(*j*) = 0 for *j* outside the interval [*n* − *N*_0_, *n* + *N*_0_].

A subset of the autocorrelation values (lags) from *Eq. 1* were used to compute a test statistic for each *v*(*n*) with the same lags (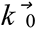 = [*k*_1_, *k*_2_, …, *k_m_*]) used for all voltage samples. Building on *Eq. 1*, for each *v*(*n*), a test statistic *c_n_* was computed as *c_n_* = 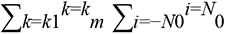 *v(n-i)v(n-i-k)* (*Eq. 2*), where *Eq. 2* is the sum of the *c_n_*(*k*) from *Eq. 1* specified by 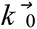. 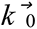 was set at 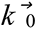 = [1, 2] because we found that these values effectively separate the distributions of the test statistics {*c_n_*} for samples of noise and samples of activity across a broad range of recording quality.

Finally, activity was considered present at *v*(*n*) only when *c_n_* was greater than a detection threshold *T. T* was set as the maximum of a set of {*c_n_*} corresponding to the {*v*(*n*)} in one contiguous second of the voltage recording where activity was confirmed to be absent (typically the first second of the recording), and was set this way for each individual voltage recording to account for differences in baseline noise levels. Fictive locomotor bursts were detected and grouped into episodes, and the burst and episode properties were determined as follows. Episode duration is the time from the onset of the first burst of an episode to the offset of the final burst in the same episode. Burst duration is the time from the onset to the offset of each burst, as defined by *c_n_* and described above. Burst frequency is the inverse of the interburst period (IBP), which is defined for each pair of bursts within episodes as the time from the onset of the first burst to the onset of the second burst. Peak burst frequency is the inverse of burst period, which is defined as the shortest time duration from onset of one burst to onset of the subsequent burst in an episode of fictive swimming. IBPs between episodes are excluded from burst frequency analysis.

### Calcium Imaging and Analysis

For calcium imaging of V3-INs, *Tg(vglut2a:Gal4ff)^nns20^;Tg(UAS:GCaMP6s)^nk13a^* zebrafish larvae were used. For calcium imaging of MNs, Calcium Green-1 AM (Life Technologies, Grand Island, NY) dye was loaded into MNs using the protocol for Rhod-2 AM dye loading described above. Larvae were transferred to a fixed-stage Olympus BX51 WI upright microscope with either a 20x / NA 0.5 or a 40x / NA 0.8 water dipping objective lens. A field of view containing *GCaMP6s*-expressing V3-INs in transgenic larvae or Calcium Green-1 AM-loaded MNs in wild type larvae was selected for calcium imaging and a simultaneous peripheral nerve (PN) recording was acquired. Calcium indicator fluorescence was excited with an E-Cite Series 120 Q lamp (Excelitas Technologies, Waltham, MA) and acquired at 20 Hz with a Retiga EXi camera and QCapture Pro 6.0 software (QImaging, Surry, BC). Data were acquired during spontaneous fictive swimming in 10-60 s durations for both V3-IN and MN trials. Multiple trials (typically 4) were acquired for each field of view, with at least five min between trials. Movement artifacts during data acquisition were corrected in Fiji by registering frames from all trials to a common reference image (Preibisch et al., 2009). Optical and electrophysiological data acquisition were synchronized by using the sync output of the stimulator to trigger acquisition of the PN recording.

For all calcium imaging experiments, raw fluorescence values were determined by drawing a region of interest around each identified V3-IN or MN soma and using the Plot Z-axis Profile command in Fiji. Fluorescence from a region of interest containing background signal was subtracted from raw soma fluorescence measurements to reduce the effect of photobleaching. Baseline fluorescence (F_0_) was defined as the mean fluorescence in the first 10 frames of the recording. To calculate ΔF/F, F_0_ was subtracted from fluorescence for each frame (F_(t)_), then divided by F_0_; ΔF/F = (F_(t)_ - F_(0)_) / F_(0)_. A neuron was considered active if ΔF/F was 10% or greater.

### Targeted Laser Ablation of V3-INs

Acute ablation of the V3-INs was performed using a Micropoint pulsed nitrogen pumped dye laser (Andor Technology, Belfast, Northern Ireland). Light (λ = 435 nm) was delivered via an Olympus BX51 WI microscope using a 60x / NA 1.0 water dipping objective lens. The laser was focused to apply maximal light intensity to the imaging plane of the microscope, the intensity of the laser was calibrated to produce a 4.5 – 5.0 µm hole in a mirror slide (this optimal intensity was determined empirically), and the x-y coordinates of the laser point in the field of view were indicated on a digital display. Three to six dpf *Tg(vglut2a:DsRed)^nns9^* zebrafish larvae were anesthetized in 0.02% Tricaine in zebrafish ringers solution and embedded laterally in a gel composed of 3% methylcellulose and 0.02% Tricaine in zebrafish ringers saline. A coverslip was placed over the gel and secured at the edges with agarose to stabilize the position of the larvae.

Targeted, bilateral laser ablation of DsRed^+^ V3-INs in the ventral spinal cord was performed by positioning the microscope objective such that a neuron was in the x-y position at which the laser was focused. Laser pulses were applied to each V3-IN soma at 33 Hz for 10 s. Laser intensity was empirically determined *a priori* such that >95% of the targeted V3-INs were not present 24 h post-ablation. Following the train of laser pulses, the microscope was repositioned to focus on another identified V3-IN, and the protocol was repeated. Larvae were embedded for approximately 50 min in order to perform bilateral ablation of ∼140 identified V3-INs between body segment (S) 5 and S20. To confirm that laser ablation of targeted V3-INs was effective and persistent, the lack of DsRed fluorescence in the lesioned spinal cord of *Tg(vglut2a:DsRed)^nns9^* larvae was confirmed ∼24 hours post-ablation. Sham larvae were treated identically to V3 ablation larvae, but the x-y position of the laser was offset by 5 – 20 µm from each V3 cell, producing non-specific damage to neurons other than the V3 population. Rather than kill a specific cell type as a sham control which might have a specific effect, non-specific laser damage was preferred.

### Statistical Analysis

Statistical analysis of data was performed using two-sample *t*-test, ANOVA test, Holm-Sidak post-hoc test, Kolmogorov-Smirnov test, and least-squares linear regression. Statistical tests were carried out using SigmaPlot 12 and 14 software (SyStat Software, San Jose, CA), Matlab (Mathworks, Natick, MA), or Microsoft Excel (Microsoft, Seattle, WA). An α level of 0.05 was used to determine statistical significance. Data are reported as means and standard deviation (Mean, SD).

### Software Accesibility

Custom Matlab code for the quantification of the distribution of V3-IN cell bodies and peripheral nerve recording analysis is available upon request.

## RESULTS

### V3-INs are Individually Identifiable in the Larval Zebrafish Spinal Cord

To characterize the functional roles of V3-INs *in vivo*, it was necessary to reliably target the neurons for recording and perturbation. While there are no published transgenic zebrafish lines that specifically label neurons expressing *sim1a* (the zebrafish V3-IN marker; (Schäfer et al., 2007)), it is possible to infer V3-IN identity from their other characteristics. V3-INs proliferate ventral to the central canal and are glutamatergic (Zhang et al., 2008; Yang et al., 2010). In zebrafish, there is >90% overlap of *vesicular glutamate transporter 2* (*vglut2a*) and *sim1a* expression in the ventral spinal cord at the embryonic stage (1.5 dpf) (Yang et al., 2010). However, it is possible that the embryonic ventral glutamatergic neurons migrate or are replaced by a non-V3 population between 1.5 dpf and the free-swimming larval stage (>3 dpf).

To test the hypothesis that the ventral DsRed+ neurons observed in 4 – 6 dpf *Tg(vglut2a:DsRed)^nns9^* larvae are V3-INs, we crossed this line to *Tg(nkx2.2a:mEGFP)^vu17^*, which marks p3 progenitor neurons (Briscoe et al., 2000; Schäfer et al., 2007) (Fig. 1A). At 4 – 6 dpf, *nkx2.2a:mEGFP* expression was restricted to the region ventral to the central canal (Fig. 1C). The p3 domain of the zebrafish spinal cord also produces Kolmer-Agduhr (KA) neurons (Yang et al., 2010), and likely other neuronal types, so it was expected that there would be more *nkx2.2a:mEGFP*-expressing neurons than *vglut2a:DsRed*-expressing neurons (Fig. 1B). Approximately 94% of *vglut2a:DsRed*-expressing neurons ventral to the central canal also expressed *nkx2.2a:mEGFP* (*n* = 6 larvae, 162 neurons), and were therefore confirmed to be V3-INs (Fig. 1D). There was not a significant difference between the dorsoventral position of the *vglut2a:DsRed*-expressing neurons that co-expressed *nkx2.2a:mEGFP* and those that did not (two-sample t-test; *t* = 1.6; *p* = 0.10), so there was not a way to reliably segregate the identified V3-INs, from those that did not express *nkx2.2a:mEGFP*. However, the overwhelming majority of the DsRed^+^ neurons ventral to the central canal were confirmed to be V3-INs. The lack of confirmed transgene overlap in the remaining cells may be due to variegated transgene expression rather than the presence of a non-V3-IN ventral glutamatergic neuronal population. Based on this experiment, we treat the ventral *vglut2a:DsRed*-expressing neurons as the V3-IN population, with the caveat that a small fraction (∼6%) may be misidentified.

**Figure 1.**
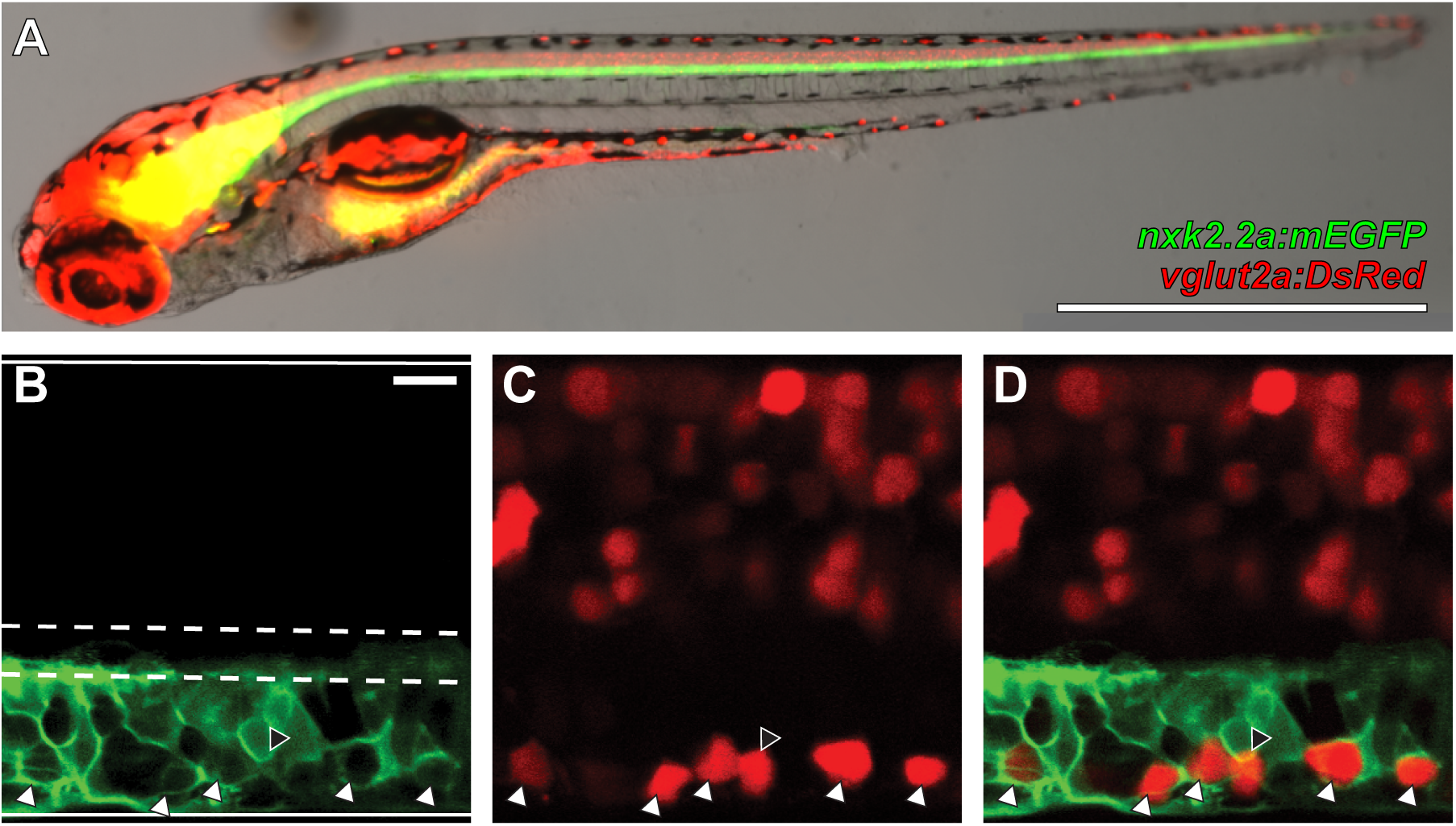
V3-INs are identified by co-localization of *vglut2a:DsRed* and *nkx2.2a:mEGFP* transgene expression in neurons located in the ventromedial spinal cord. (A) Whole-mount *Tg(nkx2.2a:mEGFP)^vu17^*;*Tg(vglut2a:DsRed)^nns9^* double transgenic zebrafish larva. (B-D) A representative single confocal optical section of the spinal cord exhibiting: (B) *nkx2.2a:mEGFP* expression, (C) *vglut2a:DsRed* expression, and (D) overlayed *nkx2.2a:mEGFP* and *vglut2a:DsRed* expression. The boundaries of the spinal cord and central canal are illustrated by solid and dashed white lines, respectively, in panel B. White triangles indicate neurons that co-express both transgenes and the black triangle indicates the only *vglut2a:DsRed* neuron ventral to the central canal that did not co-express *nkx2.2a:mEGFP* in the field of view. Scale bars: 1 mm in A and 10 µm in B-D.

### V3-INs are Located in the Ventromedial Spinal Cord and are Distributed Along the Rostrocaudal Axis of the Spinal Cord

In mice, the V3-INs migrate following differentiation to form spatially segregated dorsal and ventral populations that have different cellular properties and may be necessary for different behaviors (Borowska et al., 2013). To determine if there are multiple populations of spatially segregated V3-INs in zebrafish larvae, we used confocal microscopy to create 3-D reconstructions of the spinal cords of 4 – 6 dpf *Tg(vglut2a:DsRed)^nns9^* zebrafish larvae (Fig. 2A-B). The center of each V3-IN soma in the most rostral 2 mm of the spinal cord was tagged, and the distribution of the neurons within the spinal cord was calculated based on the imputed borders of the spinal cord (see *Methods*). Neurons in the far caudal spinal cord were not counted because the dimensions of the spinal cord taper to the extent that it was difficult to determine the location of the central canal and Mauthner axons. All V3-INs (*n* = 4 larvae; 797 neurons) were located medial of the Mauthner axon (Fig. 2C). V3-IN density was highest in the rostral spinal cord but remained close to 1 cell per 10 µm throughout the cord (Fig. 2D). The V3-IN somas were approximately 5 µm in diameter in the rostrocaudal axis, so a density of ∼1 cell per 10 µm thoroughly tiled the ventral spinal cord. The majority of V3-INs were located in the most medial and most ventral 25% of the spinal cord (Figs. 2E & F). There was no parasagittal plane through the medial spinal cord that was unoccupied by V3-INs (Fig. 2E), which suggested that the V3-IN population may be considered a midline population. However, the density of V3-INs was not highest at the sagittal midline of the spinal cord, and when cell density was plotted along the population axis, the point of highest density was adjacent to the spinal cord midline (Fig. 2G). Therefore, we propose that distinct left and right populations of V3-INs are present, but that there are not multiple ipsilateral populations of spatially segregated V3-INs in larval zebrafish.

**Figure 2.**
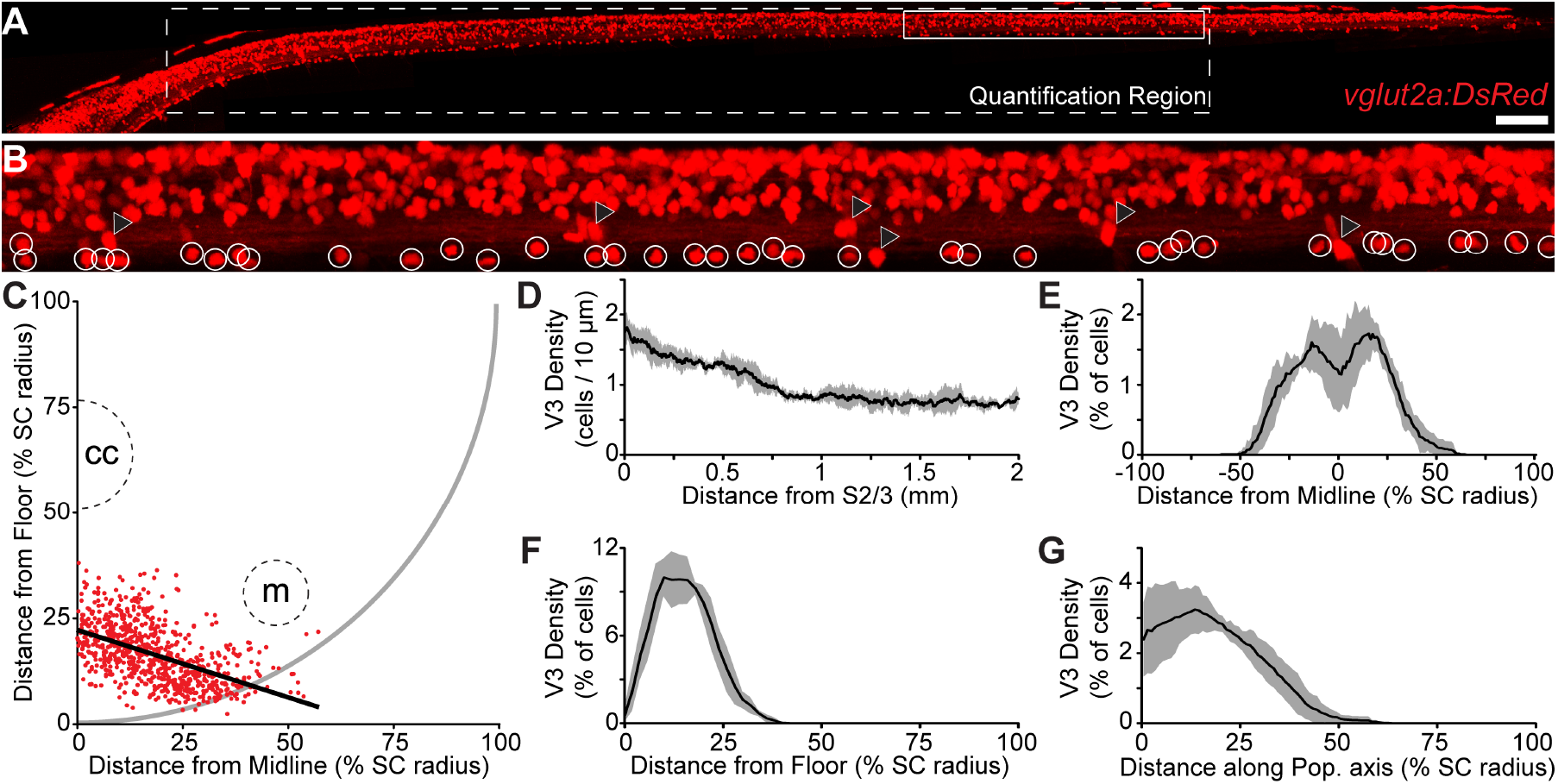
Distribution of V3-IN Cell Bodies in the Spinal Cord. (A) Maximum intensity projection of a confocal stack of the spinal cord of a 5 dpf Tg(*vglut2a:DsRed*)*^nns9^* zebrafish larva. The dashed white outline indicates the region of the cord in which cells were quantified. The solid white outline is the region of the image shown at higher magnification in (B). (B) A higher magnification field of view. White circles indicate V3 neurons, black triangles indicate DRG neurons located lateral of the spinal cord. (C) Distribution of location of all quantified V3-INs (red dots) projected in the coronal plane of the spinal cord. The schematic outline of the spinal cord (solid grey line), as well as the location of the central canal (cc) and Mauthner axon (m), are approximate. The heavy black line indicates the line of best fit that best describes the axis of the V3-IN population. (D-G) The windowed moving average of V3-IN density along the rostrocaudal axis (D; window width: 250 µm), mediolateral axis (E; window width: 15% SC radius), dorsoventral axis (F; window width: 15% SC radius), and along the population axis illustrated in panel C (G; window width: 15% SC radius). Black lines are the mean, grey area indicates the standard deviation. SC: spinal cord. Scale bar: 100 µm in A and 19 µm in B.

### The Majority of V3-INs are Active During Spontaneous Fictive Swimming

In mice, there is only indirect evidence supporting temporal correlation between V3-IN activity and locomotion (Borowska et al., 2013). To test the hypothesis that activity in the V3-INs is appropriately timed to provide excitatory drive to MNs during locomotion, we monitored the activity of the V3-INs during spontaneous fictive swimming using a genetically encoded calcium indicator (GCaMP6s; Fig. 3). The activity of V3-INs and fictive locomotor output during spontaneous swimming was measured via simultaneous calcium imaging and peripheral nerve recording using 4 - 6 dpf *Tg(vglut2a:Gal4ff)^nns20^;Tg(UAS:GCaMP6s)^nk13a^* larvae (Fig. 3B-C). We found that the majority, 45 of 48 (94%), of V3-INs were active (ΔF/F > 10%) during spontaneous fictive swimming episodes (*n* = 6 larvae, 15 trials). Therefore, this is the first direct evidence that V3-INs are active during fictive locomotion, which supports the participation of V3-INs in producing locomotor activity.

**Figure 3.**
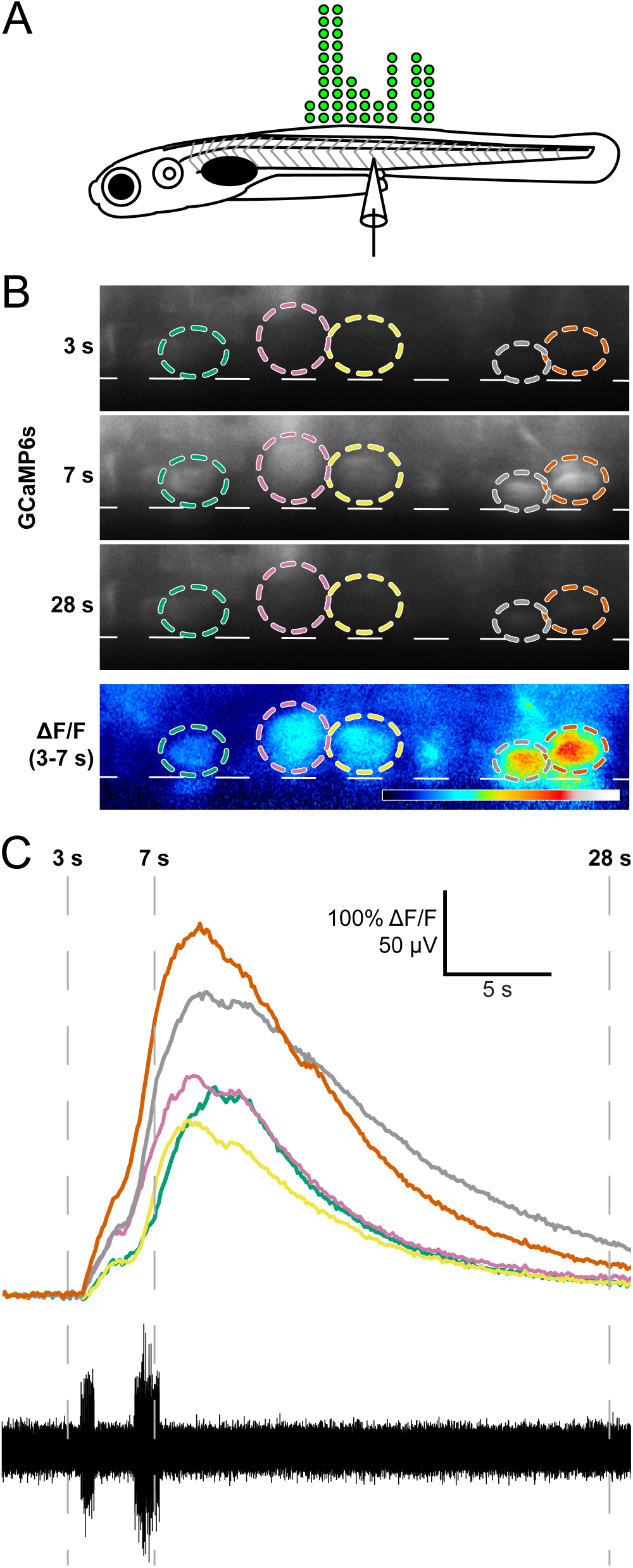
V3-IN activity is temporally correlated to fictive locomotion. (A) Schematic representing the number and distribution of V3-INs (green circles) along the rostrocaudal axis of the spinal cord monitored during calcium imaging experiments. (B) Each V3-IN in the field of view in Tg(*vglut2a:Gal4ff*)*^nns20^*;Tg(*UAS:GCaMP6s*)*^nk13a^* double transgenic larvae was assigned a ROI during spontaneous fictive swimming. Panels in (A) show raw GCaMP6s fluorescence (F) and correspond to numbered time-points in (B). Bottom panel in A shows calculated ΔF/F between 3 and 7 s time-points. Dashed lines represent ventral boundary of the spinal cord. (B) Synchronized recordings of GCaMP6s fluorescence (top) and fictive swimming (bottom) allowed identification of swimming-related neuronal activity.

### Ablation of V3-INs Does Not Affect Episodic or Burst Properties of Spontaneous Fictive Swimming

The previous experiments demonstrated that V3-INs are positioned appropriately and have neurotransmitter phenotype and activity consistent with providing excitatory drive to motor neurons; however, demonstration of the function of the V3 neurons or their necessity for behavior has not been described. To determine whether canonical features of fictive swimming (episode duration, burst (tail-beat) frequency, and burst duration) are modulated by V3-IN activity, we performed laser ablation of a large percentage of the V3-IN population (>95%; ∼140 V3-identified somas across body segments 5 to 20; Fig. 4). Three experimental groups were used: 1) The V3 ablation group, in which all V3-INs on both sides of the spinal cord between S5 and S20 were laser ablated (*n* = 6 larvae; 145 (SD 9.6) neurons per larva). Representative examples of targeted laser ablation efficacy within a restricted region (Fig. 4A-B) and spatial localization (Fig. 4C-D) are shown. 2) The sham ablation group, in which laser pulses were delivered 5 – 20 µm rostral to each identified V3-IN cell body (*n* = 6 larvae; 130 (SD 10.8) sham targets per larva). 3) The unmanipulated control group was not embedded or exposed to laser pulses (*n* = 6 larvae).

**Figure 4.**
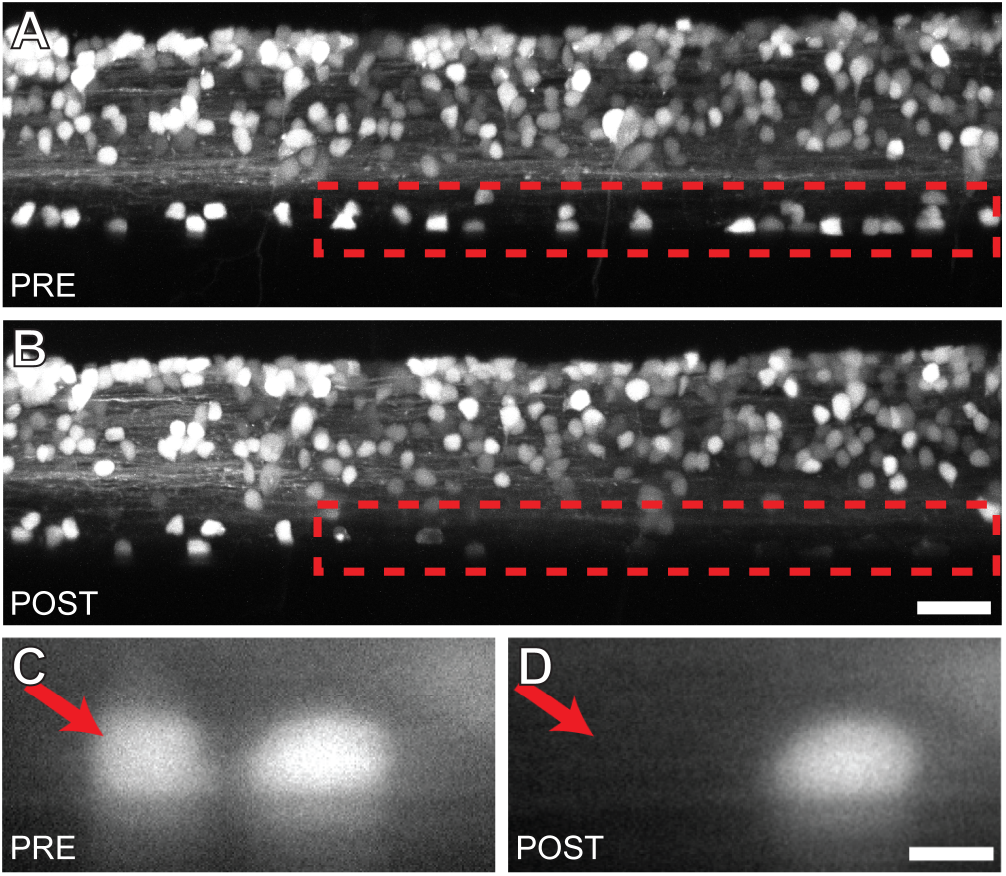
Targeted ablation of V3-INs is both efficient and spatially restricted. (A-B) Confocal stacks of the same Tg(*vglut2a:DsRed*)*^nns9^* zebrafish larva before (A) and 12 hours after (B) laser ablation of the V3-INs in the region indicated (red dashed lines). (C-D) A magnified view of two V3-INs before (left) and after (right) a single cell laser ablation. Scale bars: 20 μm in A and B and 5 μm in C and D.

The day following laser ablation or sham ablation, fictive swimming was measured using extracellular peripheral nerve recordings. Episode duration, burst frequency, and burst duration were not significantly different between unmanipulated, sham and ablation experimental groups during spontaneous swimming (one-way ANOVAs; all F < 1.3; all p > 0.30; Table 1). Further, an effect on the coordination of rostrocaudal (ipsilateral) and side-to-side (contralateral) locomotor activity was not observed (data not shown). These results indicate that the population of spinal V3-INs do not play a significant role in establishing episode duration, burst frequency (a proxy for tail-beat frequency), burst duration, or coordination during fictive locomotion in larval zebrafish.

**Table 1.**
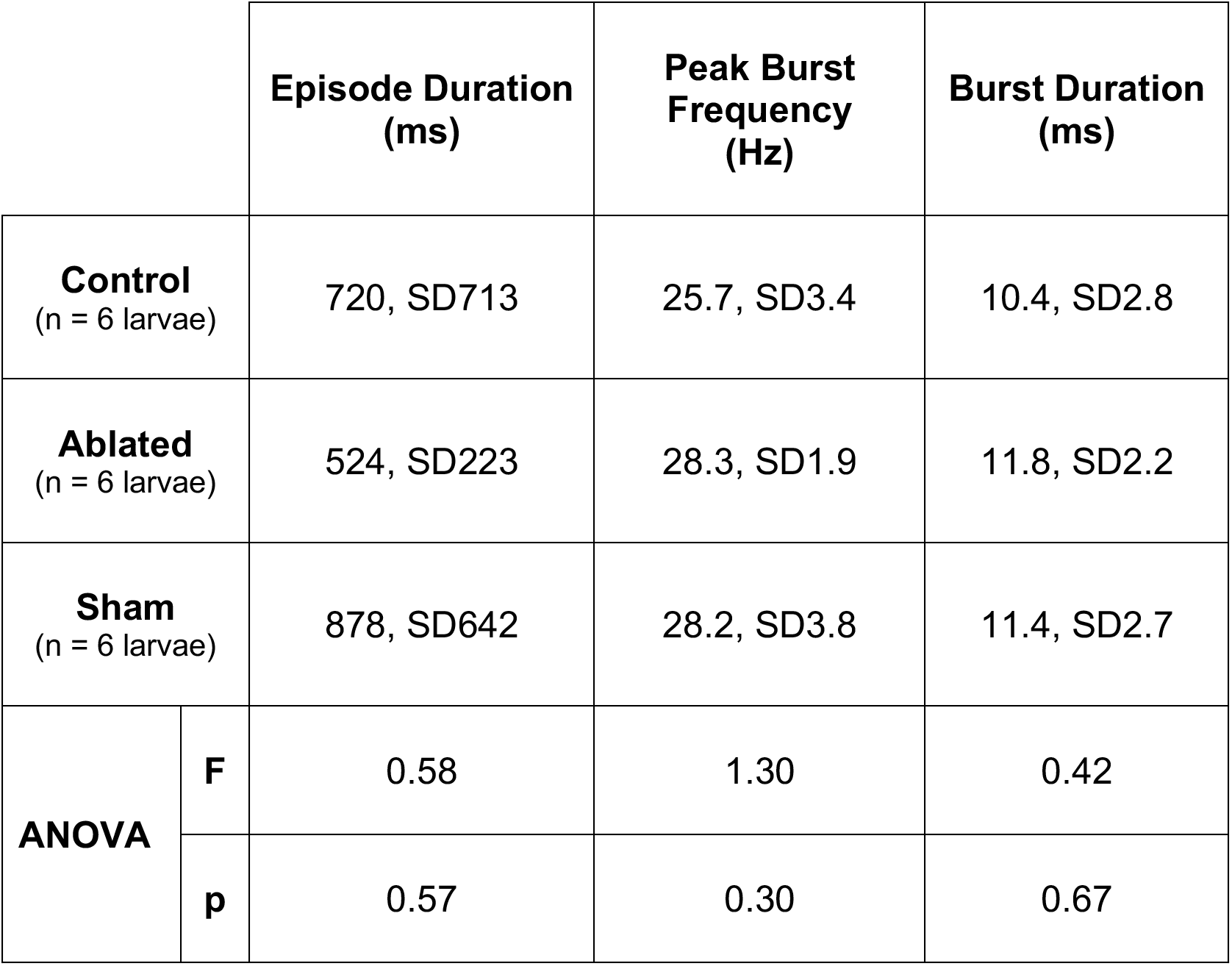
Measures of the episodic and burst properties from peripheral nerve recordings during fictive locomotion in zebrafish larvae. All values presented as mean, standard deviation.

### Ablation of V3-INs Reduces the Proportion of Active Motor Neurons but Does Not Affect Peak Locomotor Frequency During Spontaneous Fictive Swimming

Since the ablation of spinal V3-INs in zebrafish larvae did not affect episode duration, burst frequency, or burst duration (Table 1), we tested the hypothesis that ablation of V3-INs reduces locomotor amplitude by decreasing the number of active MNs during spontaneous fictive swimming. To ablate V3-INs without also photo-damaging or bleaching motor neurons, we used a two-step neuronal targeting strategy. First, V3-INs were identified and ablated using the *Tg(vglut2a:DsRed) ^nns9^* transgenic line. Following recovery from laser ablation, motor neurons were loaded with a cell-permanent calcium indicator, Calcium Green-1, AM. To verify that spinal neurons loaded with AM dye (Calcium Green-1 and/or Rhod-2) were MNs, we first confirmed motor neuronal specificity of GFP expression in the *parga^mn2Et^* enhancer trap line (Balciunas et al., 2004). *Parga^mn2Et^* larvae immunolabeled for the cholinergic marker choline acetyltransferase (ChAT) revealed that most GFP-expressing neurons co-labeled with ChAT antibodies (*n* = 5; 91.3% (SD2.1); Fig. 5A-A’’). A small percentage of dorsally-positioned ChAT+ cells did not colocalize with GFP (4.1% (SD2.8); Fig. 5A-A’’, arrowheads); these cells were considered members of a subset of cholinergic spinal interneurons that modulate MN excitability recently identified in zebrafish (Bertuzzi and Ampatzis, 2018). Thus, GFP expression in the *parga^mn2Et^* enhancer trap line was specific to MNs.

**Figure 5.**
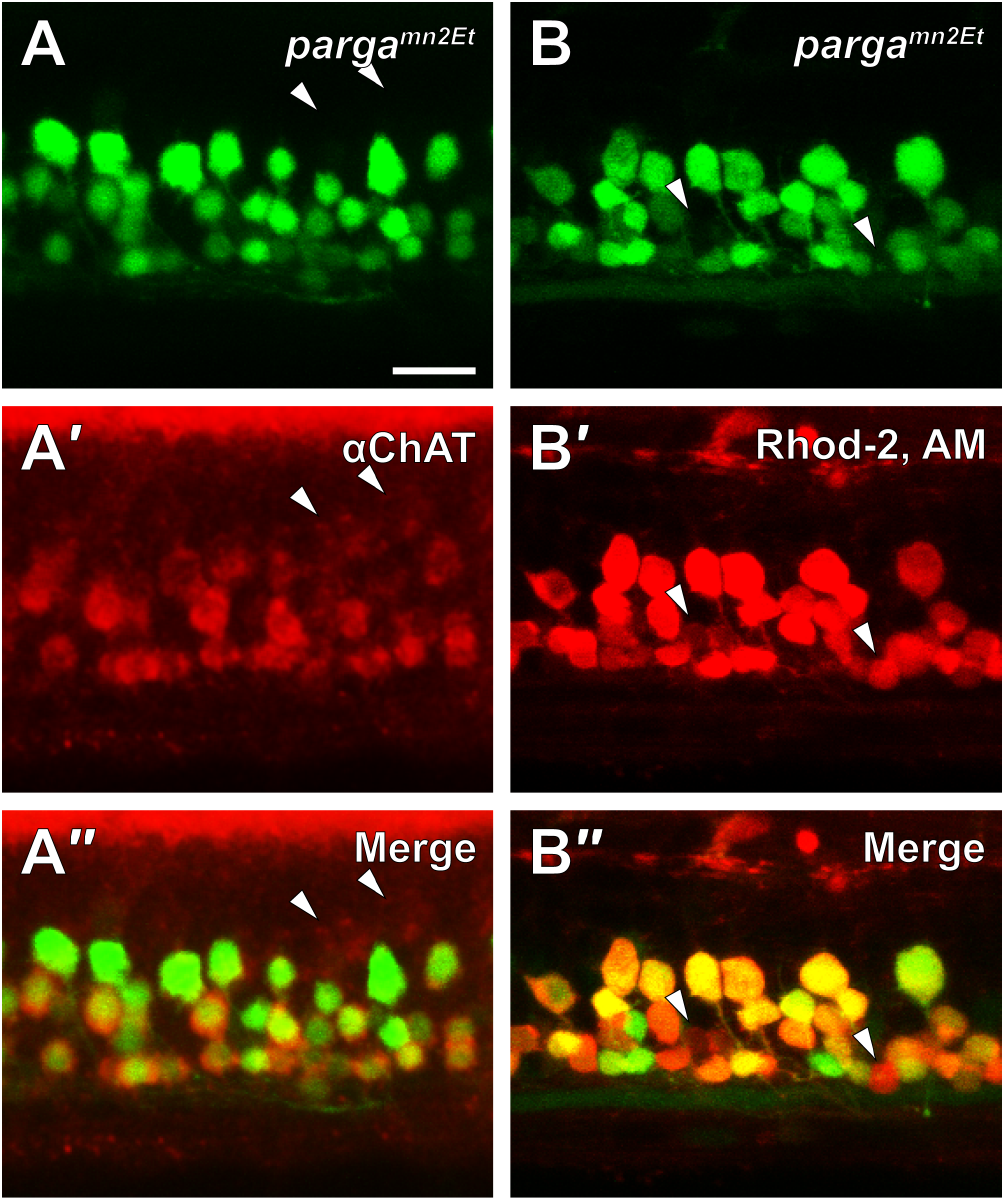
Spinal neurons loaded with rhodamine-2 AM dye are definitively identified as MNs. (A) The EGFP-expressing *parga^mn2Et^* enhancer trap line (A, green) was labeled with antibodies to choline acetyl transferase (ChAT; A′, red). Colabeling (A″) of EGFP with ChAT antibodies indicated that EGFP expression in the *parga^mn2Et^* line is a marker of spinal motor neurons. White arrowheads indicate ChAT-positive cells that do not contain EGFP (non-MNs). (B) Live *parga^mn2Et^* larvae were anesthetized, skinned, and incubated in rhodamine-2 AM dye (Rhod-2, AM). EGFP expression (B, green) and Rhod-2, AM labeling (B′, red), showed that Rhod-2, AM specifically labeled all motor neurons (B″). White arrowheads indicate two Rhod-2, AM-labeled cells that did not contain EGFP. Scale bar: 20 μm.

Next, to confirm that loading of AM dye was restricted to MNs, the skin was removed from both sides of the body from *parga^mn2Et^* larvae, in which GFP expression was shown to be limited to MNs (Fig. 5A-A’’), and MNs were labeled with Rhod-2 AM dye (*n* = 4; Fig. 5B-B’’). Rhod-2 AM, which was not used for calcium imaging experiments because it did not produce a strong calcium signal, was used as a proxy for Calcium Green-1 AM since the excitation/emission wavelengths of Calcium Green-1 AM overlapped with, and could not be disambiguated from, GFP expressed in *parga^mn2Et^* MNs. The majority of GFP-expressing MNs co-labeled with Rhod-2 AM (84.3% (SD4.3); Fig. 5B-B’’). A small percentage of GFP-negative cells contained Rhod-2 AM (5.4% (SD0.5); Fig. 6B-B’’, arrowheads). Taken together, these results confirmed that MNs were selectively labeled with AM dye and, therefore, the cell permeable calcium indicator (Calcium Green-1 AM) could be used to monitor *in vivo* calcium activity in spinal MNs.

**Figure 6.**
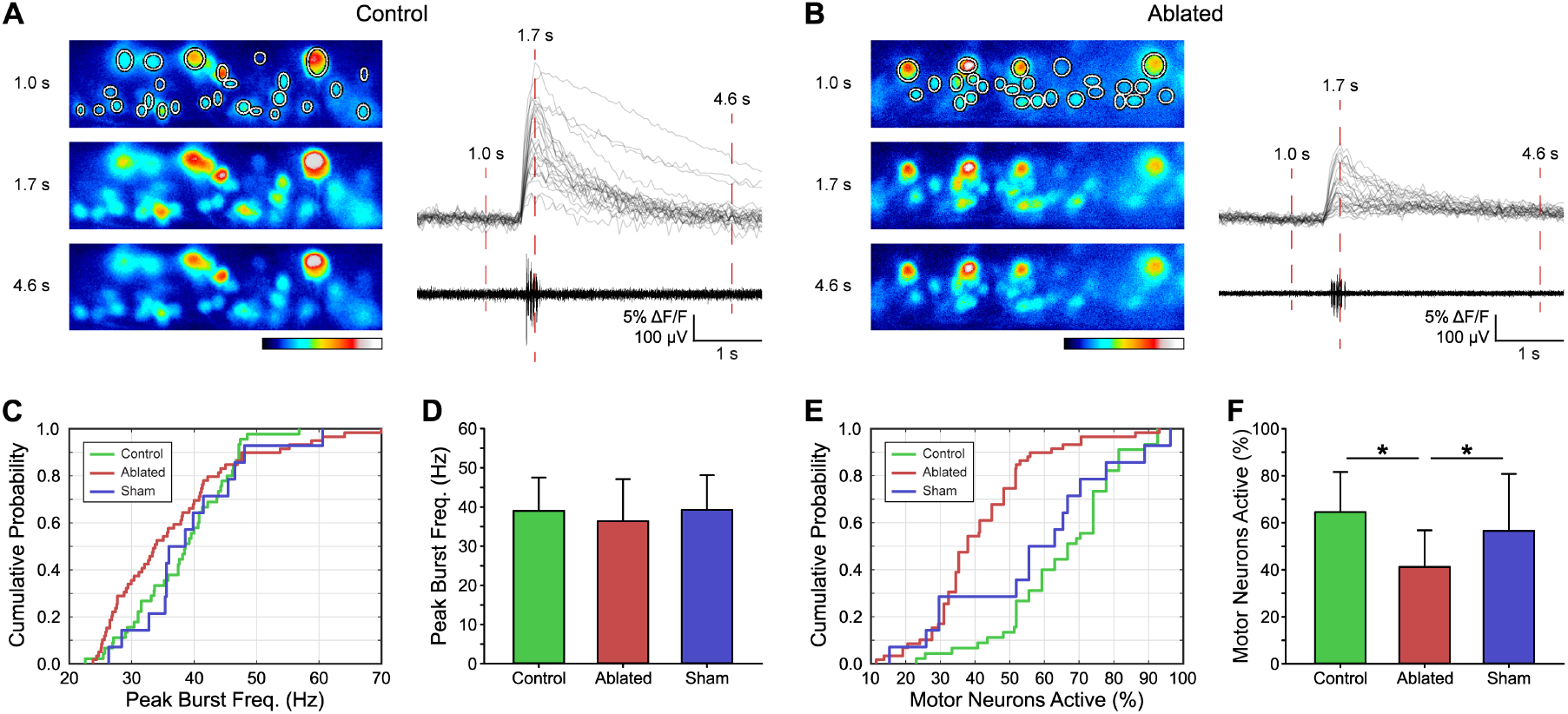
V3-IN laser ablation reduces the proportion of motor neurons that are active during fictive swimming. MNs in *Tg(vglut2a:DsRed)^nns9^* larvae were loaded with Calcium Green-1 AM dye and assigned ROIs during recording. Synchronous MN fluorescence and fictive swimming recordings were compared between Control, V3-IN Ablated, and Sham ablated preparations. (A and B) Pseudocolor Calcium Green-1 AM fluorescence panels correspond to numbered time-points in aligned ΔF/F (each gray line represents a MN) and peripheral nerve traces in a Control preparation (A) and a V3-IN Ablated preparation (B). Color indicates fluorescence intensity. (C-F) Cumulative probability distributions and mean Peak Burst Frequencies (C and D) and percent of MNs active (10% ΔF/F threshold) during fictive swimming episodes (E and F). Asterisks represent significant differences. Error bars represent SD.

To examine the effects of V3-IN ablation on the production of fictive locomotor vigor, we determined the proportion of spinal MNs that were active during spontaneous fictive swimming episodes. MN activity in *Tg(vglut2a:DsRed)^nns9^* larvae was measured using *in vivo* calcium imaging by loading MNs with Calcium Green-1 AM, while peak burst frequency during spontaneous fictive locomotor activity was measured using extracellular PN recordings (Fig. 6A-B). We quantified the proportion of active MNs and peak burst frequencies during spontaneous fictive swimming for Control (50 swim episodes, 4 larvae), Sham Ablated (14 swim episodes, 2 larvae), and V3-IN Ablated (59 swim episodes, 3 larvae) groups. Importantly, there were no differences in the properties of the fictive swimming produced by these larvae, consistent with our prior experiments (Table 1). Neither the distribution of peak burst frequencies (Kolmogorov-Smirnov test, D < 0.33, p > 0.14 for all group comparisons; Fig. 6C), nor the mean peak burst frequencies, were different between Control, Sham, and Ablation Groups (Control = 39.0 Hz (SD8.5); Sham = 36.4 Hz (SD10.7); Ablated = 39.3 Hz (SD8.9); one-way ANOVA, F = 5.0, p = 0.08; Fig. 6D).

Assessment of the probability distributions showed that the proportion of active MNs in the Ablated group was significantly reduced compared to both the Control (Kolmogorov-Smirnov test = 0.68, p < 0.001) and Sham (Kolmogorov-Smirnov test = 0.54, p = 0.001) groups (Fig. 6E). The proportion of active MNs was not significantly different between the Control and Sham groups (Kolmogorov-Smirnov test, D = 0.24, p = 0.50; Fig. 6E). Further, V3-IN ablation had a main effect of significantly reducing the proportion of active MNs during spontaneous fictive swimming (Fig. 6F; Control mean = 65.2% (SD16.7); Sham mean = 56.6% (SD24.2); Ablated mean = 41.2% (SD15.6); one-way ANOVA, F = 25.3; p < 0.001). Pairwise multiple comparisons (Holm-Sidak; Fig. 6F) revealed significant differences for the proportion of active MNs between both the Control and Ablated groups (t = 7.02, p < 0.001) and the Sham and Ablated groups (t = 2.98, p = 0.007), but significant differences were not found between Control and Sham groups (t = 1.54, p = 0.10). These results indicate that V3-INs participate in regulating locomotor amplitude by recruitment of spinal MNs but do not regulate burst frequency during spontaneous fictive locomotion.

## DISCUSSION

The goal of this study was to characterize the potential role of V3-INs in regulating locomotor amplitude in genetically tractable and optically accessible larval zebrafish. Locomotor amplitude is determined by the recruitment of motor units appropriate to the desired movement: small units for fine behaviors and large units for large movements (Zajac and Faden, 1985; Mendell, 2005). This recruitment pattern of small to large motor units, the so-called size principle, is well established in both limbed (mammalian; terrestrial locomotion) and non-limbed (swimming) locomotor systems (Henneman et al., 1965; Gustafsson and Pinter, 1984a, 1984b; Cope and Pinter, 1995; McLean et al., 2007; Gabriel et al., 2011). However, investigations into the pre-motor neural mechanisms that contribute to the regulation of locomotor amplitude have been difficult in mammals for two primary reasons. First, mammalian spinal locomotor circuits are complicated by the requirements to control and coordinate limbs and joints, rather than simply produce a smooth gradient of motion intensity (Grillner, 2003; Grillner and El Manira, 2020). Second, despite recent progress in tool development, *in vivo* functional manipulations of mammalian spinal locomotor circuits using mini-scopes and fiber-optics are both challenging and limited for long-term recordings in freely locomoting animals (Montgomery et al., 2015; Samineni et al., 2017; Canales et al., 2018). Zebrafish larvae provide several advantages over mammalian models for *in vivo* studies: the spinal cords of fish are less complicated than mammals, zebrafish are recognized as a powerful neurogenetic model system, and the translucent nature of zebrafish larvae makes them particularly appropriate for *in vivo* optical methods of investigation. In this study, we found that, unlike the mammalian spinal circuit, V3-INs in the larval zebrafish spinal cord are contained within a single, ventral anatomical domain (Fig. 1-2) and are not necessary to produce the fictive locomotor rhythm (Table 1) (Zhang et al., 2008; Danner et al., 2019). Instead, zebrafish V3-INs are swim-active neurons (Fig. 3) that contribute to the recruitment of motor neurons during fictive locomotion (Fig. 6), and therefore likely participate in the recruitment of motor units during free-moving locomotion.

### V3-INs are a population of previously uncharacterized ventral vglut2a+ neurons in larval zebrafish

Here, we report that almost all ventrally-located neurons labeled by the *Tg(vglut2a:DsRed)^nns9^* and *Tg(vglut2a:Gal4ff)^nns20^* transgenic lines are V3-IN cells (Figs. 1-4). The V3-INs, a numerous ventromedial population, are distributed across the rostrocaudal axis of the spinal cord (Fig. 2). It is unlikely that the glutamatergic V3-INs include other identified ventrally-located spinal neurons, such as the GABAergic Kolmer-Agduhr cells (Dale et al., 1987; Bernhardt et al., 1992; Montgomery et al., 2016) or the ventral serotonergic neurons (Van Raamsdonk et al., 1996; Brustein et al., 2003; McLean and Fetcho, 2004; Montgomery et al., 2018). For a more full understanding of the structural and functional relationship of the V3-INs, it will be critical to determine if the diversity of cell morphology in the V3-IN cell population is similar to the diversity of V3-INs in the mammalian spinal cord (Borowska et al., 2013; Deska-Gauthier et al., 2020). In mammals, the morphology of V3-IN subtypes is correlated with their circuit function (Borowska et al., 2013). However, we have not found evidence for differential recruitment of V3-INs during fictive swimming at various frequencies (Fig. 3), which would indicate potential functional heterogeneity. It may be that in larval zebrafish V3-INs are a primitive class of excitatory spinal interneurons, as has been previously proposed for *Engrailed1-*expressing inhibitory neurons in zebrafish (Higashijima et al., 2004). However, analysis of cell morphology in zebrafish larvae is complicated by ongoing development in the nervous system and the rostrocaudal developmental gradient (Cole and Ross, 2001). Further experiments are necessary to clarify if different morphologies represent distinct functional sub-types of V3-INs, developmental stages of a single morphological family, or regional specializations of the V3-INs.

### Zebrafish V3-INs are active during fictive locomotion

Results indicating that V3-IN activity is directly correlated to fictive locomotor activity in vertebrate preparations has not been convincingly presented. In mice, there is indirect evidence indicating temporal correlation between V3-IN activity and locomotion (Borowska et al., 2013). Through simultaneous optical and electrophysiological recordings, we showed that activity in most V3-INs is correlated to fictive swimming (Fig. 3) and is therefore appropriately timed to provide excitatory drive to MNs during locomotion. Thus, this is the first direct evidence in any vertebrate preparation that V3-INs are active during fictive locomotion, which provides supporting evidence that they may participate in locomotor activity.

### V3-IN activity is necessary for motor neuron recruitment, but not necessary for rhythm generation

Given that V3-INs have been shown to affect the regularity and coordination of locomotor activity in mice (Zhang et al., 2008), it was surprising that laser ablation of V3-INs in larval zebrafish had no effect on episode duration, mean burst frequency, or burst duration during fictive swimming in larval zebrafish (Table 1). One interpretation of these findings is that despite similarities in gene expression and neurotransmitter profile, premotor interneuron function is divergent between axial and tetrapod locomotion. The latter argument has been advanced regarding the V2a-INs (Dougherty and Kiehn, 2010; Dougherty et al., 2013); the different effects (or lack thereof) of V3-IN ablation in zebrafish and mice may be a second example of the trend. An important difference between the current study and mouse knock-out experiments is the mechanism of ablation. In the current study, V3-INs were ablated acutely, and little time (∼24 hours) was given for circuit compensation. In the mouse experiments, on the other hand, V3-INs were ablated or silenced chronically (Zhang et al., 2008; Rabe Bernhardt et al., 2012), leaving opportunity for physiological compensation. Acute silencing experiments using optogenetics could be performed in both species (Arrenberg et al., 2009; Hägglund et al., 2013), and may demonstrate that this difference between species is due to the experimental manipulation rather than the underlying neural circuit.

Finally, a subset of identified V2a excitatory spinal interneurons has been shown to control the speed of locomotion in a variety of vertebrates (Zhong et al., 2011; Ausborn et al., 2012; Ampatzis et al., 2014; Gosgnach et al., 2017). Of particular interest, a recent study also ascribed a role for regulating locomotor amplitude/vigor to a morphologically and functionally distinct subset of the V2a-INs (type II, non-bursting) in larval and adult zebrafish (Song et al., 2018; Menelaou and McLean, 2019). Taken together, these results indicate that the general population of V2a-INs is involved in regulating both locomotor speed and amplitude. To date, however, a population of spinal interneurons that selectively regulates locomotor amplitude has not been identified. Our work provides evidence that, in zebrafish larvae, V3-INs contribute selectively to locomotor amplitude. Specifically, we show that targeted ablation of V3-INs (Fig. 4) reduces the proportion of active motor neurons during fictive swimming (Fig. 6E-F) but does not affect the range of locomotor frequencies produced (Fig. 6C-D). Thus, we propose that V3-INs are a source of excitation in the vertebrate locomotor neural circuitry that regulates the amplitude of locomotor output independently of locomotor frequency.

### Conclusion

The work presented here addressed an important concept regarding motor control; specifically, a limited understanding of the cellular properties that produce locomotor amplitude in vertebrates. We showed, directly for the first time, that V3-INs in zebrafish larvae are active during *in vivo* fictive locomotion (Fig. 3) and contribute to motor neuron recruitment. Importantly, targeted ablation of the V3-IN spinal cord population does not affect locomotor frequency (speed; Fig. 6C-D), which clarifies their role in motor control (amplitude) rather than rhythm generation (frequency). Thus, we propose that the V3-IN population regulates locomotor amplitude independently of frequency, which is an important functional difference between the V3-INs and the general population of V2a-INs since the latter regulate both frequency and amplitude (Zhong et al., 2011; Ausborn et al., 2012; Ampatzis et al., 2014; Gosgnach et al., 2017; Song et al., 2018; Menelaou and McLean, 2019).

## Author Contributions

TDW, JEM and MAM Designed the Research; TDW, JEM, AJB, JHP and MAM Performed the Research; TDW and MAM Wrote the Paper

## Acknowledgements

The authors appreciate the helpful suggestions and feedback from Drs. Ronald Calabrese and Ronald Harris-Warrick. We also thank Kayce Vanpelt for technical support and Marc Tye and the staff of the University of Minnesota Zebrafish Core Facility for animal care. Finally, we thank Drs. Koichi Kawakami and Misha Ahrens for kindly providing *Tg(UAS:GCaMP6s)^nk13a^* fish. Current affiliation for TDW is Vizgen, Cambridge, MA 02138 and AJB is Florida State University, Department of Psychology, Tallahassee, FL 32306

## REFERENCES

Ampatzis K, Song J, Ausborn J, El Manira A (2014) Separate microcircuit modules of distinct v2a interneurons and motoneurons control the speed of locomotion. Neuron 83:934–943 Available at: http://linkinghub.elsevier.com/retrieve/pii/S0896627314006291 [Accessed October 19, 2017].

Arrenberg AB, Del Bene F, Baier H (2009) Optical control of zebrafish behavior with halorhodopsin. Proc Natl Acad Sci U S A 106:17968–17973 Available at: http://www.pnas.org/cgi/doi/10.1073/pnas.0906252106 [Accessed October 23, 2017].

Ausborn J, Mahmood R, El Manira A (2012) Decoding the rules of recruitment of excitatory interneurons in the adult zebrafish locomotor network. Proc Natl Acad Sci U S A 109:E3631–9 Available at: http://www.pnas.org/cgi/doi/10.1073/pnas.1216256110 [Accessed October 19, 2017].

Bainbridge R (1958) The Speed of Swimming of Fish as Related to Size and to the Frequency and Amplitude of the Tail Beat. J Exp Biol 35.

Balciunas D, Davidson AE, Sivasubbu S, Hermanson SB, Welle Z, Ekker SC (2004) Enhancer trapping in zebrafish using the Sleeping Beauty transposon. BMC Genomics 5.

Bellardita C, Kiehn O (2015) Phenotypic Characterization of Speed-Associated Gait Changes in Mice Reveals Modular Organization of Locomotor Networks. Curr Biol 25:1426–1436 Available at: http://www.ncbi.nlm.nih.gov/pubmed/25959968 [Accessed October 23, 2017].

Bernhardt RR, Patel CK, Wilson SW, Kuwada JY (1992) Axonal trajectories and distribution of GABAergic spinal neurons in wildtype and mutant zebrafish lacking floor plate cells. J Comp Neurol 326:263–272.

Bertuzzi M, Ampatzis K (2018) Spinal cholinergic interneurons differentially control motoneuron excitability and alter the locomotor network operational range. Sci Rep 8.

Borowska J, Jones CT, Zhang H, Blacklaws J, Goulding M, Zhang Y (2013) Functional subpopulations of V3 interneurons in the mature mouse spinal cord. J Neurosci 33:18553–18565 Available at: http://www.jneurosci.org/cgi/doi/10.1523/JNEUROSCI.2005-13.2013 [Accessed October 23, 2017].

Briscoe J, Pierani A, Jessell TM, Ericson J (2000) A homeodomain protein code specifies progenitor cell identity and neuronal fate in the ventral neural tube. Cell 101:435–445 Available at: http://www.ncbi.nlm.nih.gov/pubmed/10830170 [Accessed October 22, 2017].

Brustein E, Chong M, Holmqvist B, Drapeau P (2003) Serotonin Patterns Locomotor Network Activity in the Developing Zebrafish by Modulating Quiescent Periods. J Neurobiol.

Canales A, Park S, Kilias A, Anikeeva P (2018) Multifunctional Fibers as Tools for Neuroscience and Neuroengineering. Acc Chem Res 51.

Chopek JW, Nascimento F, Beato M, Brownstone RM, Zhang Y (2018) Sub-populations of Spinal V3 Interneurons Form Focal Modules of Layered Pre-motor Microcircuits. Cell Rep 25.

Cole LK, Ross LS (2001) Apoptosis in the developing zebrafish embryo. Dev Biol 240:123–142.

Cope T, Pinter M (1995) The Size Principle: Still Working After All These Years. Physiology 10:280–286.

Crone SA, Zhong G, Harris-Warrick R, Sharma K (2009) In mice lacking V2a interneurons, gait depends on speed of locomotion. J Neurosci 29:7098–7109 Available at: http://www.jneurosci.org/cgi/doi/10.1523/JNEUROSCI.1206-09.2009 [Accessed October 23, 2017].

Dale N, Roberts A, Ottersen OP, Storm-Mathisen J (1987) The morphology and distribution of “Kolmer-Agduhr cells”, a class of cerebrospinal-fluid-contacting neurons revealed in the frog embryo spinal cord by GABA immunocytochemistry. Proc R Soc Lond B Biol Sci.

Danner SM, Zhang H, Shevtsova NA, Borowska-Fielding J, Deska-Gauthier D, Rybak IA, Zhang Y (2019) Spinal V3 Interneurons and Left–Right Coordination in Mammalian Locomotion. Front Cell Neurosci.

Deska-Gauthier D, Borowska-Fielding J, Jones CT, Zhang Y (2020) The temporal neurogenesis patterning of spinal p3–V3 interneurons into divergent subpopulation assemblies. J Neurosci 40.

Donley JM, Dickson KA (2000) Swimming kinematics of juvenile kawakawa tuna (Euthynnus affinis) and chub mackerel (Scomber japonicus). J Exp Biol 203:3103–3116 Available at: http://www.ncbi.nlm.nih.gov/pubmed/11003821 [Accessed October 23, 2017].

Dougherty KJ, Kiehn O (2010) Firing and Cellular Properties of V2a Interneurons in the Rodent Spinal Cord. J Neurosci 30:24–37 Available at: http://www.jneurosci.org/cgi/doi/10.1523/JNEUROSCI.4821-09.2010.

Dougherty KJ, Zagoraiou L, Satoh D, Rozani I, Doobar S, Arber S, Jessell TM, Kiehn O (2013) Locomotor Rhythm Generation Linked to the Output of Spinal Shox2 Excitatory Interneurons. Neuron 80:920–933.

Eklöf-Ljunggren E, Haupt S, Ausborn J, Dehnisch I, Uhlén P, Higashijima S, El Manira A (2012) Origin of excitation underlying locomotion in the spinal circuit of zebrafish. Proc Natl Acad Sci U S A 109:5511–5516 Available at: http://www.pnas.org/cgi/doi/10.1073/pnas.1115377109 [Accessed October 19, 2017].

Ericson J, Rashbass P, Schedl A, Brenner-Morton S, Kawakami A, van Heyningen V, Jessell TM, Briscoe J (1997) Pax6 controls progenitor cell identity and neuronal fate in response to graded Shh signaling. Cell 90:169–180 Available at: http://www.ncbi.nlm.nih.gov/pubmed/9230312 [Accessed October 22, 2017].

Gabriel JP, Ausborn J, Ampatzis K, Mahmood R, Eklöf-Ljunggren E, El Manira A (2011) Principles governing recruitment of motoneurons during swimming in zebrafish. Nat Neurosci 14:93–99 Available at: http://www.nature.com/doifinder/10.1038/nn.2704 [Accessed October 19, 2017].

Gosgnach S, Bikoff JB, Dougherty KJ, El Manira A, Lanuza GM, Zhang Y (2017) Delineating the diversity of spinal interneurons in locomotor circuits. J Neurosci 37.

Grillner S (2003) The motor infrastructure: From ion channels to neuronal networks. Nat Rev Neurosci 4.

Grillner S, El Manira A (2020) Current principles of motor control, with special reference to vertebrate locomotion. Physiol Rev 100.

Grillner S, Jessell TM (2009) Measured motion: searching for simplicity in spinal locomotor networks. Curr Opin Neurobiol 19:572–586 Available at: http://www.ncbi.nlm.nih.gov/pubmed/19896834 [Accessed October 20, 2017].

Gustafsson B, Pinter MJ (1984a) An investigation of threshold properties among cat spinal alpha-motoneurones. J Physiol 357:453–483.

Gustafsson B, Pinter MJ (1984b) Relations among passive electrical properties of lumbar alpha-motoneurones of the cat. J Physiol 356:401–431.

Hägglund M, Dougherty KJ, Borgius L, Itohara S, Iwasato T, Kiehn O (2013) Optogenetic dissection reveals multiple rhythmogenic modules underlying locomotion. Proc Natl Acad Sci U S A 110:11589–11594.

Heglund NC, Taylor CR, Mcmahon TA (1974) Scaling stride frequency and gait to animal size: Mice to horses. Science (80- ) 186.

Henneman E, Somjen G, Carpenter DO (1965) Functional Significance of Cell Size in Spinal Motoneurons. J Neurophysiol 28:560–580.

Higashijima S, Masino MA, Mandel G, Fetcho JR (2004) Engrailed-1 expression marks a primitive class of inhibitory spinal interneuron. J Neurosci 24:5827–5839 Available at: http://www.ncbi.nlm.nih.gov/pubmed/15215305 [Accessed October 19, 2017].

Hunter JR, Zweifel JR (1971) Swimming speed, tail beat frequency, tail beat amplitude, and size in jack makerel, Trachurus symmetricus, and other fishes. Fish Bull 69:253–267 Available at: http://swfsc.noaa.gov/publications/CR/1971/7114.PDF [Accessed October 23, 2017].

Jayne BC, Lauder GV (1995) Speed effects on midline kinematics during steady undulatory swimming of largemouth bass, Micropterus salmoides. J Exp Biol 198:585–602 Available at: http://www.ncbi.nlm.nih.gov/pubmed/9318295 [Accessed October 23, 2017].

Kiehn O (2011) Development and functional organization of spinal locomotor circuits. Curr Opin Neurobiol 21:100–109 Available at: http://www.ncbi.nlm.nih.gov/pubmed/20889331 [Accessed October 20, 2017].

Kimura Y, Satou C, Fujioka S, Shoji W, Umeda K, Ishizuka T, Yawo H, Higashijima S (2013) Hindbrain V2a Neurons in the Excitation of Spinal Locomotor Circuits during Zebrafish Swimming. Curr Biol 23:843–849 Available at: http://www.ncbi.nlm.nih.gov/pubmed/23623549 [Accessed October 20, 2017].

Kirby BB, Takada N, Latimer AJ, Shin J, Carney TJ, Kelsh RN, Appel B (2006) In vivo time-lapse imaging shows dynamic oligodendrocyte progenitor behavior during zebrafish development. Nat Neurosci.

Lin S, Li Y, Lucas-Osma AM, Hari K, Stephens MJ, Singla R, Heckman CJ, Zhang Y, Fouad K, Fenrich KK, Bennett DJ (2019) Locomotor-related V3 interneurons initiate and coordinate muscles spasms after spinal cord injury. J Neurophysiol.

Ljunggren EE, Haupt S, Ausborn J, Ampatzis K, El Manira A (2014) Optogenetic activation of excitatory premotor interneurons is sufficient to generate coordinated locomotor activity in larval zebrafish. J Neurosci 34:134–139 Available at: http://www.jneurosci.org/cgi/doi/10.1523/JNEUROSCI.4087-13.2014 [Accessed October 19, 2017].

McLean DL, Fan J, Higashijima S, Hale ME, Fetcho JR (2007) A topographic map of recruitment in spinal cord. Nature 446:71–75 Available at: http://www.ncbi.nlm.nih.gov/pubmed/17330042 [Accessed October 19, 2017].

McLean DL, Fetcho JR (2004) Ontogeny and innervation patterns of dopaminergic, noradrenergic, and serotonergic neurons in larval zebrafish. J Comp Neurol 480:38–56 Available at: http://www.ncbi.nlm.nih.gov/pubmed/15515022 [Accessed October 19, 2017].

Mendell LM (2005) The size principle: A rule describing the recruitment of motoneurons. J Neurophysiol 93:3024–3026.

Menelaou E, McLean DL (2019) Hierarchical control of locomotion by distinct types of spinal V2a interneurons in zebrafish. Nat Commun 10.

Menelaou E, VanDunk C, McLean DL (2014) Differences in the morphology of spinal V2a neurons reflect their recruitment order during swimming in larval zebrafish. J Comp Neurol 522:1232–1248 Available at: http://doi.wiley.com/10.1002/cne.23465 [Accessed October 19, 2017].

Miyasaka N, Morimoto K, Tsubokawa T, Higashijima S, Okamoto H, Yoshihara Y (2009) From the olfactory bulb to higher brain centers: genetic visualization of secondary olfactory pathways in zebrafish. J Neurosci 29:4756–4767 Available at: http://www.jneurosci.org/cgi/doi/10.1523/JNEUROSCI.0118-09.2009 [Accessed October 20, 2017].

Montgomery JE, Wahlstrom-Helgren S, Wiggin TD, Corwin BM, Lillesaar C, Masino MA (2018) Intraspinal serotonergic signaling suppresses locomotor activity in larval zebrafish. Dev Neurobiol.

Montgomery JE, Wiggin TD, Rivera-Perez LM, Lillesaar C, Masino MA (2016) Intraspinal serotonergic neurons consist of two, temporally distinct populations in developing zebrafish. Dev Neurobiol 76:673–687 Available at: http://doi.wiley.com/10.1002/dneu.22352 [Accessed October 19, 2017].

Montgomery KL, Yeh AJ, Ho JS, Tsao V, Iyer SM, Grosenick L, Ferenczi EA, Tanabe Y, Deisseroth K, Delp SL, Poon ASY (2015) Wirelessly powered, fully internal optogenetics for brain, spinal and peripheral circuits in mice. Nat Methods 12.

Müller UK, van Leeuwen JL (2004) Swimming of larval zebrafish: ontogeny of body waves and implications for locomotory development. J Exp Biol 207:853–868 Available at: http://www.ncbi.nlm.nih.gov/pubmed/14747416 [Accessed October 22, 2017].

Muto A, Lal P, Ailani D, Abe G, Itoh M, Kawakami K (2017) Activation of the hypothalamic feeding centre upon visual prey detection. Nat Commun.

Pierani A, Brenner-Morton S, Chiang C, Jessell TM (1999) A sonic hedgehog-independent, retinoid-activated pathway of neurogenesis in the ventral spinal cord. Cell 97:903–915 Available at: http://www.ncbi.nlm.nih.gov/pubmed/10399918 [Accessed October 22, 2017].

Preibisch S, Saalfeld S, Tomancak P (2009) Globally optimal stitching of tiled 3D microscopic image acquisitions. Bioinformatics.

Rabe Bernhardt N, Memic F, Gezelius H, Thiebes A-L, Vallstedt A, Kullander K (2012) DCC mediated axon guidance of spinal interneurons is essential for normal locomotor central pattern generator function. Dev Biol 366:279–289 Available at: http://linkinghub.elsevier.com/retrieve/pii/S0012160612001807 [Accessed October 20, 2017].

Samineni VK et al. (2017) Fully implantable, battery-free wireless optoelectronic devices for spinal optogenetics. Pain 158.

Satou C, Kimura Y, Hirata H, Suster ML, Kawakami K, Higashijima S -i. (2013) Transgenic tools to characterize neuronal properties of discrete populations of zebrafish neurons. Development 140:3927–3931 Available at: http://www.ncbi.nlm.nih.gov/pubmed/23946442 [Accessed October 20, 2017].

Schäfer M, Kinzel D, Winkler C (2007) Discontinuous organization and specification of the lateral floor plate in zebrafish. Dev Biol 301:117–129 Available at: http://linkinghub.elsevier.com/retrieve/pii/S001216060601219X [Accessed October 25, 2017].

Schindelin J, Arg I, Arganda-Carreras I, a-Carreras, Frise E, Kaynig V, Longair M, Pietzsch T, Preibisch S, Rueden C, Saalfeld S, Schmid B, Tinevez J-Y, White D, Hartenstein V, Eliceiri K, Tomancak P, Cardona A (2012) Fiji: an open-source platform for biological-image analysis. Nat Methods 9.

Song J, Dahlberg E, El Manira A (2018) V2a interneuron diversity tailors spinal circuit organization to control the vigor of locomotor movements. Nat Commun.

Tytell ED (2004) The hydrodynamics of eel swimming II. Effect of swimming speed. J Exp Biol 207:3265–3279 Available at: http://www.ncbi.nlm.nih.gov/pubmed/15326203 [Accessed October 23, 2017].

Van Raamsdonk W, Bosch TJ, Smit-Onel MJ, Maslam S (1996) Organisation of the zebrafish spinal cord: distribution of motoneuron dendrites and 5-HT containing cells. Eur J Morphol.

Videler JJ, Müller UK, Stamhuis EJ (1999) Aquatic vertebrate locomotion: Wakes from body waves. J Exp Biol 202.

Wardle, Videler, Altringham (1995) Tuning in to fish swimming waves: body form, swimming mode and muscle function. J Exp Biol 198.

Webb PW (1975) Hydrodynamics and energetics of fish propulsion. Bull Fish Res Board Canada 190:1–159 Available at: http://www.dfo-mpo.gc.ca/library/1486.pdf [Accessed October 23, 2017].

Webb PW, Kostecki PT, Stevens ED (1984) The Effect of Size and Swimming Speed on Locomotor Kinematics of Rainbow Trout. J Exp Biol 109.

Wiggin TD, Anderson TM, Eian J, Peck JH, Masino MA (2012) Episodic swimming in the larval zebrafish is generated by a spatially distributed spinal network with modular functional organization. J Neurophysiol 108:925–934 Available at: http://jn.physiology.org/cgi/doi/10.1152/jn.00233.2012 [Accessed October 19, 2017].

Yang L, Rastegar S, Strähle U (2010) Regulatory interactions specifying Kolmer-Agduhr interneurons. Development 137:2713–2722 Available at: http://dev.biologists.org/cgi/doi/10.1242/dev.048470 [Accessed October 20, 2017].

Zajac FE, Faden JS (1985) Relationship among recruitment order, axonal conduction velocity, and muscle-unit properties of type-identified motor units in cat plantaris muscle. J Neurophysiol 53:1303–1322.

Zhang Y, Narayan S, Geiman E, Lanuza GM, Velasquez T, Shanks B, Akay T, Dyck J, Pearson K, Gosgnach S, Fan CM, Goulding M (2008) V3 Spinal Neurons Establish a Robust and Balanced Locomotor Rhythm during Walking. Neuron 60:84–96 Available at: http://linkinghub.elsevier.com/retrieve/pii/S0896627308008064 [Accessed October 20, 2017].

Zhong G, Sharma K, Harris-Warrick RM (2011) Frequency-dependent recruitment of V2a interneurons during fictive locomotion in the mouse spinal cord. Nat Commun 2:274 Available at: http://www.nature.com/doifinder/10.1038/ncomms1276 [Accessed October 19, 2017].

